# ConvDip: A convolutional neural network for better EEG Source Imaging

**DOI:** 10.1101/2020.04.09.033506

**Authors:** Lukas Hecker, Rebekka Rupprecht, Ludger Tebartz van Elst, Jürgen Kornmeier

## Abstract

The EEG is a well-established non-invasive method in neuroscientific research and clinical diagnostics. It provides a high temporal but low spatial resolution of brain activity. In order to gain insight about the spatial dynamics of the EEG one has to solve the inverse problem, i.e. finding the neural sources that give rise to the recorded EEG activity. The inverse problem is ill-posed, which means that more than one configuration of neural sources can evoke one and the same distribution of EEG activity on the scalp. Artificial neural networks have been previously used successfully to find either one or two dipoles sources. These approaches, however, have never solved the inverse problem in a distributed dipole model with more than two dipole sources. We present ConvDip, a novel convolutional neural network (CNN) architecture that solves the EEG inverse problem in a distributed dipole model based on simulated EEG data. We show that (1) ConvDip learned to produce inverse solutions from a single time point of EEG data and (2) outperforms state-of-the-art methods on all focused performance measures. It is more flexible when dealing with varying number of sources, produces less ghost sources and misses less real sources than the comparison methods. It produces plausible inverse solutions for real EEG recordings from human participants. (4) The trained network needs less than 40 ms for a single prediction. Our results qualify ConvDip as an efficient and easy-to-apply novel method for source localization in EEG data, with high relevance for clinical applications, e.g. in epileptology and real time applications.

## 2 Introduction

### 2.1 The EEG and the inverse problem

Electroencephalography (EEG) is among the most used imaging techniques for noninvasive measurements of electromagnetic brain activity. Its main advantage over other methods (e.g. functional magnetic resonance imaging; fMRI) is the high temporal resolution, which comes at the cost of a considerably low spatial resolution (Luck, 2014). As a consequence, EEG was mainly used to study temporal brain dynamics at fine time scales. In the past decades, however, there has been a steady growth of interest in the neural sources underlying the EEG signal (Grech et al., 2008; He, Sohrabpour, Brown, & Liu, 2018; Koles, 1998; Michel & Brunet, 2019; R. D. Pascual-Marqui, 1999). The non-invasive estimations of neural generators, based on their projections to the scalp electrodes/sensors, constitutes an inverse problem. Without further constraints it is ill posed because it lacks an unique solution, since multiple configurations of neural sources can produce identical topographies of electrical activity at the scalp (see e.g. Nunez & Srinivasan, 2006).

Invasive multimodal methods (e.g. using combined EEG and electrocorticogram) help to bridge the gap between scalp recordings and neural generators and thus in handling the inverse problem in this constellation. However, access to these methods is limited and the conclusions that can be drawn are constrained by various factors such as placement of the cortical electrodes or coverage of restricted brain areas that project to the scalp electrodes. Combined EEG-fMRI has also been shown as a useful tool in providing insight into the spatiotemporal dynamics of the EEG (Ritter & Villringer, 2006). However, the costs of this technique are considerably high and the relation between electromagnetic and metabolic dynamics is yet not fully understood.

### 2.2 Classical approaches to the EEG inverse problem

Introducing some constraints on the solution, one can solve the inverse problem or at least reduce the number of possible solutions. One approach is the equivalent current dipole model, which is based on the assumption that the source of a signal, measured with the EEG, can be modeled by a single (or sometimes few) dipole(s) Delorme, Palmer, Onton, Oostenveld, and Makeig (2012); Kavanagk, Darcey, Lehmann, and Fender (1978); Scherg (1990). Although single-dipole sized sources are too small to generate detectable scalp potentials (Nunez & Srinivasan, 2006) they can produce reasonable results (Ebersole, 1994; Lantz, Holub, Ryding, & Rośen, 1996; Sharma et al., 2018; Willemse, Hillebrand, Ronner, Peter Vandertop, & Stam, 2016).

A more physiologically realistic approach is the distributed dipole model in which activity is expected to extend over larger areas of the brain (as opposed to tiny dipoles). Distributed dipole models aim to find the 3D distribution of neural activity underlying the EEG measurement (Hämäläinen & Ilmoniemi, 1994; R. Pascual-Marqui, Michel, & Lehmann, 1994). A distributed dipole model proposes that sources of the EEG are better modeled using hundreds to many thousand dipoles and therefore aim to find a distributed activity that can explain the EEG data. This model can be viewed opposed to single- to few-dipole models, which assume that EEG data can be sufficiently modeled using point sources.

A popular family of distributed dipole solutions is the *Minimum Norm Estimates* (MNE, Hämäläinen & Ilmoniemi, 1994; Ioannides, Bolton, & Clarke, 1990), which aim to find the source configuration that minimizes the required power to generate a given potential at the scalp electrodes. Low Resolution Electromagnetic Tomography (LORETA) is a famous proponent of the MNE-family that assumes sources to be smoothly distributed (R. Pascual-Marqui et al., 1994). In the most sophisticated version, exact LORETA (eLORETA, R. D. Pascual-Marqui, 2007) showed zero localization error when localizing single sources in simulated data (R. D. Pascual-Marqui, 2007).

Another popular family of inverse solutions is the beamforming approach. The linear constrained minimum variance (LCMV) beamforming approach is a spatial filter that assumes that neural sources are uncorrelated and in which portions of the data that don’t belong to the signal are suppressed (Van Veen, van Drongelen, Yuchtman, & Suzuki, 1997). Drawbacks of the LCMV approach are the susceptibility to imprecisions in the forward model and that correlated sources are often not found.

Growing interest towards a bayesian notation of the inverse problem could be observed in the past two decades (Chowdhury, Lina, Kobayashi, & Grova, 2013a; Friston et al., 2008; Wipf & Nagarajan, 2009). One prominent bayesian approach is the maximum entropy on the mean (MEM) method, which aims to make the least assumptions on the current distribution by maximizing entropy (Amblard, Lapalme, & Lina, 2004; Chowdhury, Lina, Kobayashi, & Grova, 2013b; Grova et al., 2006).

### 2.3 Artificial neural networks (ANN) and inverse solutions

ANN based inverse solutions follow a data-driven approach and were of increasing interest in the past years (Awan, Saleem, & Kiran, 2019). A large number of simulated EEG data samples is used to train an ANN to correctly map electrode-space signals to source-space locations (Jin, McCann, Froustey, & Unser, 2017). Long training periods are usually required for an ANN to generalize beyond the training set. After successful training it is capable of predicting the coordinates and orientations of a dipole correctly, given only the measurements at the scalp electrodes without further priors (Abeyratne, Zhang, & Saratchandran, 2001; Zhang, Yuasa, Nagashino, & Kinouchi, 1998).

Robert, Gaudy, and Limoge (2002) reviewed the literature on ANN-based methods to solve the inverse problem of single dipoles and found that all reports achieved localization errors of less than 5%. The low computational time (once trained) and robustness to measurement noise was highlighted when compared to classical approaches. However, the classical approaches were capable of achieving zero localization error in single-dipole simulations under zero-noise conditions (Hoey et al., 2000).

In an effort to predict source locations of two dipoles, Yuasa, Zhang, Nagashino, and Kinouchi (1998) presented a feed-forward network with two hidden layers and achieved localization errors in a range between 3 – 9%. They found that if multiple simulated dipoles have a sufficient distance among each other the localization success of ANNs even for multiple simulated sources is equal to classical approaches, making a strong case for the feasibility of ANNs for the EEG inverse problem, already in 1998. Although ANNs as such gained popularity in the past decade on other areas (e.g. image classification or natural language processing), the idea of ANN-based solutions to the EEG inverse problem received little further attention.

Only very recently several studies about ANN-based solutions to the inverse problem have been published. Cui et al. (2019) showed that a neural network can be trained to reconstruct the position and time course of a single source using a long-short term memory recurrent neural network (LSTM) architecture (Hochreiter & Schmidhuber, 1997). LSTMs allow to not only use single time instances of (e.g. EEG) data but instead learn from temporally lagged information.

In a recent work Razorenova et al. (2020) showed that the inverse problem for cortical potential imaging could be solved using a deep U-Net architecture, once more providing a strong case for the feasibility of ANNs to solve the EEG inverse problem (Fedorov, Koshev, & Dylov, 2020).

Tankelevich (2019) showed that a deep feed-forward network can find the correct set of source clusters that produced a given scalp signal. To our knowledge this was the first ANN approach to calculate distributed dipole solutions.

In the very recent years, convolutional neural networks (CNNs) have proven to be a useful tool in a steadily increasing number of domains, like image classification (Krizhevsky, Sutskever, & Hinton, 2012), natural language processing (Kim, 2014) and decoding of single trial EEG (Schirrmeister et al., 2017). CNNs are capable of learning patterns in data with a preserved temporal (e.g. time sequences) or spatial (e.g. images) structure by optimizing filter kernels that are convolved with a given input. Two famous CNNs are AlexNet (Krizhevsky et al., 2012) and VGG16 (Simonyan & Zisserman, 2014), that won the ImageNet classification competition in 2012 and 2014, respectively.

In the current study we explored the feasibility of CNNs to solve the EEG inverse problem. Specifically, we constructed a CNN, named *ConvDip*, that is capable of detecting multiple sources using training data with biologically plausible constraints. ConvDip solves the EEG inverse problem using a distributed dipole solution in a data-driven approach. ConvDip was trained to work on single time-instances of EEG data and predicts the position of sources from potentials measured with scalp electrodes.

## 3 Methods

A Python package to create a forward model, simulate data, train an ANN and perform predictions is available at https://github.com/LukeTheHecker/ESINet.

### 3.1 Forward model

In order to simulate EEG data realistically one has to solve the forward problem and construct a generative model (GM). An anatomical template was used as provided by the Freesurfer image analysis suite (http://surfer.nmr.mgh.harvard.edu/) called *fsaverage* (Fischl, Sereno, Tootell, & Dale, 1999).

We calculated the three shell boundary element method (BEM, Fuchs, Kastner, Wagner, Hawes, & Ebersole, 2002) head model with 5120 vertices per shell using the python package MNE (v 19, Gramfort et al., 2013) and the functions *make bem model* and *make bem solution* therein. The conductivity was set to 0.3*S/m*^2^ for brain and scalp tissue and 0.06*S/m*^2^ for the skull.

The source model was created with *p* = 5124 dipoles along the cortical surface provided by Freesurfer with icosahedral spacing. We chose *q* = 31 electrodes based on the 10-20 system (Fp1, F3, F7, FT9, FC5, FC1, C3, T7, TP9, CP5, CP1, Pz, P3, P7, O1, Oz, O2, P4, P8, TP10, CP6, CP2, Cz, C4, T8, FT10, FC6, FC2, F4, F8, Fp2). The leadfield **K** *∈* ℛ*^q×p^* was then calculated using the head model with dipole orientations fixed orthogonal to the cortical surface (see e.g. Michel & Brunet, 2019 for explanation).

Additionally, we created another forward model which we refer to as the alternative generative model (AGM). The AGM will be used to simulate data for the model evaluation and serves the purpose to avoid the *inverse crime*. The inverse crime is committed when the same generative model is used to both simulate data *and* calculate inverse solutions, leading to overoptimistic results (Wirgin, 2004). First, electrode positions were changed by adding random normal distributed noise *𝒩* (0, 2*mm*) to the X, Y and Z coordinate of each electrode. This resulted in an average displacement per electrode of *≈* 3.7*mm*. Furthermore, we changed the tissue conductivity in the BEM solution from to 0.332*S/m*^2^ for brain and scalp tissue and 0.0113*S/m*^2^ for skull tissue (values were adapted as reported in Wolters, Lew, MacLeod, & Hämäläinen, 2010). This step was done in order to introduce some alterations in the volume conduction of the brain-electric signals, which are never known precisely *a priori*. Importantly, this changes the skull-to-brain conductivity ratio from 1:50 to 1:25. Last, we changed the spacing of the dipoles along the cortical sheet from icosahedral to octahedral with a higher resolution. This resulted in 8196 dipoles in the source model of the AGM, all of which were placed at slightly different locations on the cortex compared to the source model of the GM. The idea of increasing the resolution of the source model to avoid the inverse crime was adapted from Kaipio and Somersalo (2006). All simulations in the evaluation section were carried out using this alternative generative model (AGM), whereas the calculation of the inverse solutions of cMEM, eLORETA and LCMV beamformer were based on the GM. Likewise, data used for training ConvDip was simulated using only the GM.

### 3.2 Simulations

Training data for ConvDip was created using the GM, whereas the data for the evaluation of the model was created using the AGM. Since AGM and GM have different source models we implemented a function which transforms a source vector **j***_AGM_* to a source vector **j***_GM_*. This was achieved by K nearest-neighbor interpolation, i.e. each value in the target space was assigned the average value of the K nearest neighbors in the initial space. K was set to 5 neighbors. Since the source model of AGM contained almost twice as many dipoles as that of the GM a source vector translated from AGM to GM will effectively loose total source power. We therefore normalized the resulting source vector **j***_GM_* to have equal energy (sum of all dipole moments) as **j***_AGM_*.

In summary, different sets of simulated EEG samples were generated to 1) create training data for ConvDip and 2) for evaluation of ConvDip and other inverse solutions.

Each simulation contained at least one dipole cluster which can be considered as a smooth region of brain activity. A dipole cluster was generated by selecting a random dipole in the cortical source model and then adapting a region growing approach as described in Chowdhury et al. (2013b). In brief, we recursively included all surrounding neighbors starting from a single seeding location, thereby creating a larger source extent with each iteration. The number of iterations define the neighborhood order *s*, where the first order *s*_1_ entails only the single selected dipole. Each seed location was assigned a dipole moment between 5 and 10*nAm*. The neighboring dipoles were assigned attenuated moments based on the distance to seeding location. The attenuation followed a gaussian distribution with a mean of the seeding dipole moment and a standard deviation of half the radius of the dipole cluster, yielding smooth source patches.

After generating this spatial pattern we added a temporal dimension to the data as follows. We considered an epoch length of 1 second at a sampling frequency of 100 Hz (i.e. *t* = 100 time points). The time course was modelled by a central peak of 100 ms temporal extension using half a period of a 5*Hz* sinusoidal, surrounded by zeros. Each dipole moment was then multiplied by this time course. The simulated source **J** *∈* ℛ*^p×t^* of *p* = 5124 dipoles was then projected to *q* = 31 EEG electrodes **M** *∈* ℛ*^q×t^* by solving the forward problem using the leadfield **K** *∈* ℛ*^q×p^*:

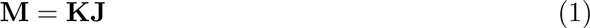

In order to generate realistic training data we added real noise from pre-existing EEG recordings, conducted with the same set of electrodes as described above. Therefore, from a set of different raw EEG recordings we extracted 200 random segments. These EEG segments were filtered beforehand using a band-pass between 0.1 and 30 Hz. Each segment contained 1 second of EEG data. The sampling frequency was reduced from 1000 Hz to 100 Hz. Baseline correction was applied based on the first 100 ms and the resulting data was re-referenced to common average. For each sample we then created 20 EEG trials by adding a randomly selected noise segment to the simulated EEG. Prior to this, the noise was scaled to yield a signal-to-noise ratio (SNR) of 1 within single trials. The average of these 20 trials, i.e. the event-related potential, should therefore exhibit a theoretical SNR of 4.5 at the ERP peak.

The extent of the sources was defined between two (*s*_2_) and five (*s*_5_) neighborhood orders, which corresponds to diameters between *≈* 21 mm and *≈* 58 mm (or 19 to 91 dipoles).

#### 3.2.1 Training data

For training ConvDip we simulated in total 100,000 samples using the GM as described above. Each sample contained between 1 and 5 source clusters of extent between 2 and 5 neighborhood orders. Since ConvDip operates on single time instances of EEG or ERP data, we only used the EEG at the signal peak as input.

#### 3.2.2 Evaluation data

Two separate sets of simulations of 1000 samples each were created for the evaluation. The first set contained samples of single source clusters with varying extents from two to seven neighborhood orders, which we will refer to as single-source set. The second set contained samples with varying numbers of source clusters from 1 to 10 and with varying extents from 2 to 7 neighborhood orders or diameters from *≈* 21*mm* and *≈* 58*mm* (or 16 to 113 dipoles in the AGM). We will refer to this as the multi-source set. Source extents have different diameters for different source models, especially in our case where the source model of GM has roughly half the number of dipoles compared to AGM. This explains why *s*_7_ for the AGM is the close to *s*_6_ in the GM. Note, that both sets were created using simulations of brain-electric activity based on an alternative generative model that differs from that used in the inversion process. This was done in order to avoid the *inverse crime* (for details see Methods section 3.1).

### 3.3 ConvDip

In this section the architecture of ConvDip, a convolutional neural network which solves the inverse problem, is described and a mathematical description in the classical notation of inverse solutions is given.

#### 3.3.1 I/O of ConvDip

Since ConvDip was designed to operate on single time instances of EEG data we extracted from each simulated data sample only the time point of the peak source activity. The EEG data at this time frame was then interpolated on a 2D image of size 7*x*11. This is necessary, given that the architecture of ConvDip (Fig. 1) requires a 2D matrix input to perform spatial convolutions on. Note, that this interpolation procedure does not add information to the EEG data. The output of ConvDip is a vector of size 5124 corresponding to the dipoles in the source model (see Fig. 1).

**Figure 1:**
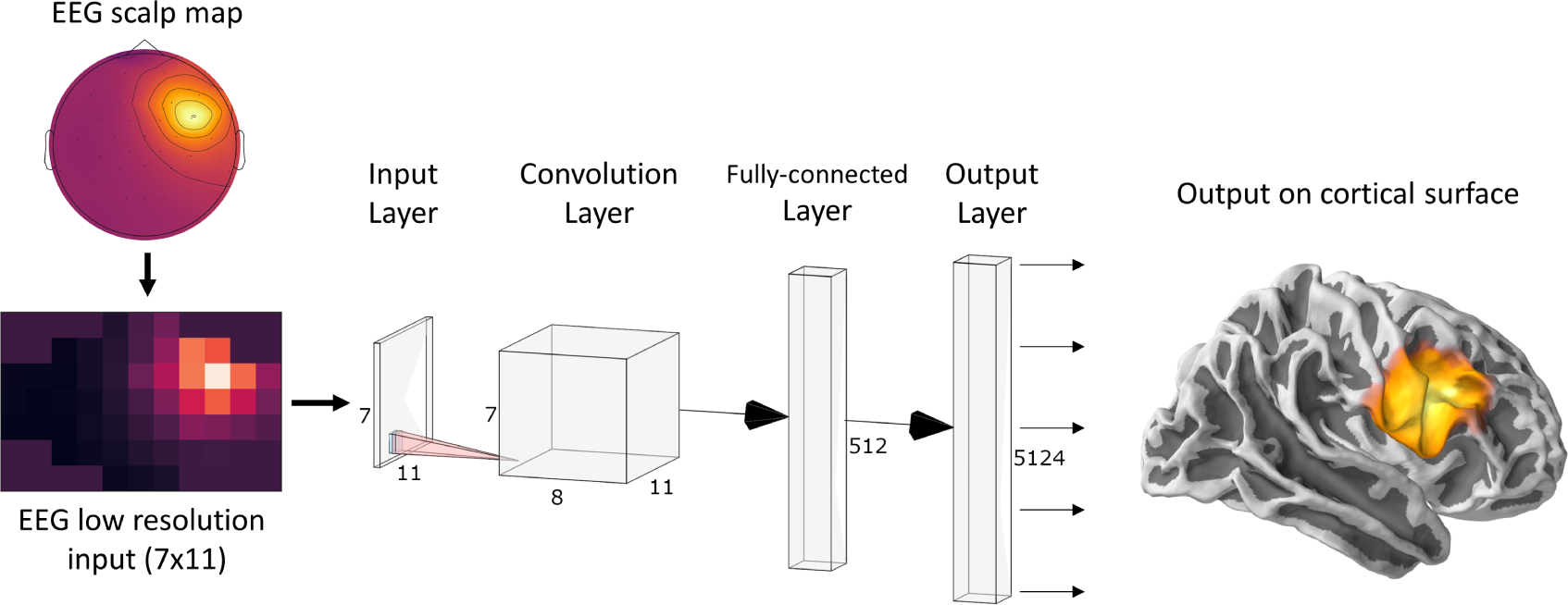
ConvDip architecture. The values from a single time point of EEG data were interpolated to get a 7×11 matrix as an input (see illustration on the bottom left). The subsequent convolution layer has only 8 convolution kernels of size 3×3 pixels. The convolution layer is followed by a fully-connected (FC) layer consisting of 512 neurons. Finally, the output layer contains 5124 neurons that correspond to the voxels in the brain (plotted on a cortical surface on the right). (This diagram was created using a web application at http://alexlenail.me/NN-SVG/)

#### 3.3.2 ConvDip architecture

The design and training of the neural network was accomplished using the Tensorflow (Abadi et al., 2016) and Keras (Chollet et al., 2015) libraries, which are based on Python 3. Training was partially accomplished on a Nvidia Titan V graphical processing unit (GPU) and an Nvidia RTX 2070.

The architecture of ConvDip (Fig. 1) is inspired by CNNs for image classification, in which the input layer is forwarded to a series of convolutional layers followed by a series of fully-connected layers followed by the output layer (e.g. AlexNet; Krizhevsky et al., 2012). In typical CNNs, the convolution layers are followed by pooling layers to reduce dimensionality and to promote invariance to an object’s position Deru and Ndiaye (2019). Notice, that in ConvDip, no pooling layers were used since spatial information would get lost or at least blurred. While invariance to the position of an object is desirable in image classification tasks (what-task), it would be detrimental in the case of source localization problems (where-task). Finally, we decided for a moderately shallow architecture with a single convolution layer followed by a fully-connected layer and an output layer.

We decided to use Rectified Linear Units (ReLUs) as activation functions after each layer (Glorot, Bordes, & Bengio, 2011; Nair & Hinton, 2010). ReLUs have shown to exhibit the best performance in our preliminary tests compared to alternatives (e.g. sigmoid function). After each of the inner layers we added a batch normalization layer (Ioffe & Szegedy, 2015) in order to speed up training and to reduce internal covariance shifts. In the current version of ConvDip the output layer consists of 5124 neurons, which equals the number of voxels in the modeled brain. ReLU activation functions were used in the output layer. Typical CNNs for regression use linear activation functions, however, since predictions are by definition non-negative in our application, ReLUs appeared to us as an appropriate alternative.

ConvDip can be described as a function 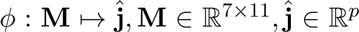, with *p* = 5124 that maps a single time instance of 2D-interpolated EEG data **M** to a source vector 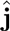:

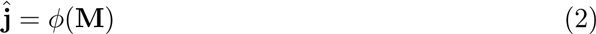

The architecture of ConvDip starts with a Convolution Layer with only 8 filters **F***_i_, i* = *{*1*, …*8*}* of size 3 *×* 3. The weights of these filters are learnt during the training period. In the forward pass the padded input EEG data **M** is convolved with each filter **F***_i_* resulting in one feature map **G***_i_ ∈* R^7^*^×^*^11^.

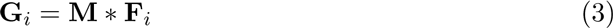

The feature maps **G***_i_, i ∈ {*1*, …,* 8*}* are stacked to a tensor **G** *∈* ℛ^7^*^×^*^11^*^×^*^8^ and reshaped to a vector 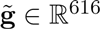 (also called *flattening*) to enable a connection to the next FC Layer. The flattened vector 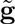 consists of 616 output nodes. Each node is connected to every neuron of the following FC Layer.

For each of the 512 neurons in the hidden fully-connected layer we transform its input 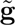 using the weight vector **w**, bias *b* and the activation function *h*:

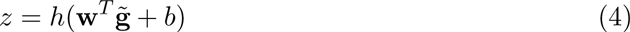

The hidden layer is finally connected to the Output Layer of 5124 neurons.

#### 3.3.3 Optimization and loss function

Convolution filters, weights and biases were optimized using adaptive momentum estimation (ADAM, Kingma & Ba, 2014) with default settings as suggested by the authors (*learning rate* = 0.001, *β*_1_ = 0.9, *β*_2_ = 0.999, *E* = 10^8^).

We tried out various loss functions for the training of ConvDip. The *mean squared error* loss is a classical loss for regression tasks. For EEG inverse solutions, however, it is not appropriate since it operates pixel-wise, i.e. the error does not provide information on how close a wrong prediction is to the true source. This is especially problematic when ConvDip is designed to find focal sources. Various approaches exist to tackle this issue, e.g. by solving an *optimal transport problem* (e.g. using Wasserstein metric) or by calculating the distance between two sets of coordinates (e.g. Hausdorff distance). We decided for the weighted Hausdorff distance (WHD) as described by Ribera, Guera, Chen, and Delp (2019)^1^.

The WHD has shown to perform well on image segmentation tasks and allowed for a fast convergence of ConvDip compared to MSE. Note, that WHD requires a normalization of the prediction and target to ensure all values are between 0 and 1. Therefore, only positional information is regarded in this loss. The rationale for this is to disentangle the problem of estimating absolute dipole moments and that of estimating dipole positions.

As mentioned above, ConvDip predicts source locations without correct global scaling (see also 2). In order to obtain the true amplitude of the sources we used Brent’s method (Brent, 1971) to find a scalar *ŝ* that minimizes the mean squared error between the forward solution (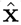) of the predicted sources 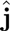) and the unscaled EEG vector **x**:

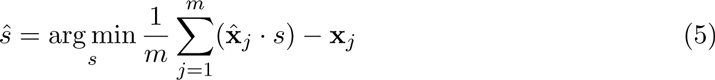

The scalar *ŝ* can then be used to scale the prediction.

### 3.4 Implementation of cMEM, eLORETA & LCMV

In order to evaluate ConvDip we calculated inverse solutions on the same set of simulations using eLORETA, LCMV beamformer and cMEM and compared the different methods with each other.

eLORETA and LCMV were carried out by the implementations in the Python library MNE. Using the same head model as described in chapter 3.1. Each inverse algorithm was subjected to each sample of the evaluation set as described in 3.2. The first 400 ms of each trial were used to estimate the noise covariance matrices. Regularization of the noise covariance was established using the standard MNE procedure as described in Gramfort et al. (2014). We choose the regularization parameter for eLORETA inverse solutions at 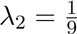 and for LCMV beamforming we set the data covariance regularization to *λ* = 0.05, which both correspond to MNE’s default parameters. Dipole orientations were restricted to be fixed orthogonal to the cortical surface.

cMEM inverse solutions were calculated with Brainstorm (Tadel, Baillet, Mosher, Pantazis, & Leahy, 2011), which is documented and freely available for download online under the GNU general public license (http://neuroimage.usc.edu/brainstorm). The same template brain (fsaverage) was used to calculate a forward model in Brainstorm. Furthermore, the BEM solution was calculated using the same parameters as in MNE. cMEM inverse solutions were calculated using a neighborhood order of 4 with temporally stable clusters. Precomputed noise covariance matrices were imported from MNE Python to ensure that inverse solutions were calculated under the same conditions in Brainstorm and MNE. After calculating all inverse solutions the files were exported again for further evaluation in Python.

### 3.5 Evaluation metrics

We calculated a number of performance metrics to assess the quality of an inverse solution. Note, that for each sample in the evaluation sets we only analyzed the central peak activity.

1. **Area under the ROC curve**: We calculated the area under the receiver operator curve (ROC) to assess the ability to estimate the correct spatial extent of sources. We adapted a similar procedure as described in Chowdhury et al. (2013a); Grova et al. (2006) with minor changes to the procedure. The dipole clusters in the target source vector **j** have a gaussian distribution of dipole moments in our simulations. Therefore, we had to set all members of all dipole clusters to unit amplitude, thereby normalizing the target source vector between 0 and 1. The estimated source vector 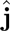 was normalized between 0 and 1 by division of the maximum. The AUC metric requires equal as many positives as there are negatives in the data. However, in our simulations only few dipoles were active (positives) compared to those not being active (negatives). Therefore, we adapted the procedure as described by Chowdhury et al. (2013a) and Grova et al. (2006) by calculating two types of AUC: Both AUCs included all positives and only differ in the selection of negatives. *AUC_close_* contained negative examples that closely surrounded the positive voxels by sampling randomly from the closest 20% voxels. This metric therefore captures how well the source extent is captured. *AUC_far_* on the other hand sampled the negatives from voxels far away from true sources, therefore capturing possible false positives in the estimated source vector 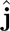. Far negatives were sampled from the 50% of the farthest negatives to the next active dipole. The overall AUC was then calculated by taking the average of *AUC_close_* and *AUC_far_*.
2. **Mean localization error (MLE)**: The Euclidean distance between the locations of the predicted source maximum and the target source maximum is a common measure of inverse solution accuracy as it captures the ability to localize the source center accurately. MLE was calculated between the positions of the maxima of 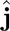 and **j**. This metric is only suitable for calculating MLE when a single source patch is present. For multiple sources we adapted the following procedure. First, we identified local maxima in both the true source vector **j** and the estimated source vector 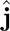. First, local maxima were identified where a voxel value is larger than all of its neighboring voxels. This then yields many local maxima which had to be filtered in order to achieve reasonable results. First, we removed all maxima whose neighbors were not sufficiently active (*<* 20% of the maximum). This takes care of false positive maxima that do not constitute an active cluster of voxels. Then, we removed those maxima that had a larger maximum voxel in close neighborhood within a radius of 30 mm. These procedures result in a list of coordinates of maxima for both **j** and 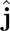. We then calculated the pairwise Euclidean distances and between the two lists of coordinates of maxima. For each true source we identified the closest estimated source and calculated the MLE by averaging of these minimum distances. We further labeled those estimated sources that were *≥* 30*mm* away from the next true maximum as *ghost sources*. True maxima that did have an estimated source within a radius of 30*mm* were labeled as *found sources*, whereas those true maxima that did not have an estimated maximum within a radius of 30*mm* were labeled as missed sources. Finally, we calculated the *percentage of found sources*, i.e. the ratio of the number of correctly identified sources and the number of all present sources.
3. **Mean squared error (MSE)**: The MSE is calculated voxel-wise between the true source vector **j** and the predicted source vector 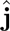:

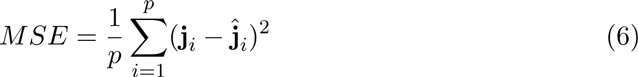
4. **Normalized mean squared error (nMSE)**: The nMSE is calculated by first nor-malizing both 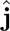 and **j** to values between 0 and 1. Then, the voxel-wise MSE is calculated as described above.

### 3.6 Statistics

Statistical comparison between the outcomes of different inverse solutions was done using an unpaired permutation test with 10^6^ permutations. The rationale behind this choice is that most distributions did not meet the criteria for parametric and rank-based tests. Additionally, Cohens *d* is given for each statistical test (Cohen, 1992).

### 3.7 Evaluation with real data

In order to evaluate the performance of ConvDip and the other inverse algorithms in a more realistic set-up we used data of a real EEG recording. The data was recorded while a single participant viewed fast presentations of faces and scrambled faces. The participant had to indicate whether a face was presented by button press. The duration of a stimulus presentation was 600 ms, followed by 200 ms of black screen. 300 trials of both faces and scrambled faces were presented, which corresponds to a total of 4 minutes.

The EEG was recorded with 31 electrodes of the 10-20 system using the ActiCap electrode cap and the Brain Vision amplifier ActiCHamp. Data was sampled at 1000*Hz* and filtered online using a band-pass of 0.01 to 120 Hz. The EEG data was imported to MNE Python and filtered using a band-pass filter between 0.1 and 30 Hz. Data was re-referenced to common average. The trials in which faces were presented were then selected and baseline corrected by subtracting the average amplitude in the interval −0.06 to 0.04 s relative to stimulus onset.

## 4 Results

In this section we evaluate the performance of ConvDip and compare it to state-of-the-art inverse algorithms, namely eLORETA, LCMV beamformer and cMEM. Note, that the evaluation set was not part of the training set of ConvDip; hence it is unknown to the model. Furthermore, all samples in the evaluation set were created using the AGM in order to avoid the inverse crime. Exemplary and representative samples are shown in Appendix A and Appendix B.

### 4.1 Evaluation with single source set

#### 4.1.1 Evaluation of source extents

We will now evaluate the ability of ConvDip to estimate the correct size of sources and to correctly localize sources with varying depth. For this purpose we used the data samples from the single-source set. An exemplary and representative sample is shown in 6.

One of the advantages of inverse algorithms such as cMEM over minimum norm solutions is their capability to estimate not only the position of a source cluster but also its spatial extent. This can be tested using the AUC metric. Chowdhury et al. (2013a) suggested that an AUC of 80% and above can be considered acceptable.

This criterion is met by all inverse algorithms in the present evaluation (Table 1). ConvDip achieves the best scores in this comparison with an overall AUC of 98%, closely followed by LCMV (95%) and cMEM (91%). Furthermore, ConvDip inverse solutions show the highest global similarity with nMSE at 0.0033, which is less than half of the error yielded by cMEM. Moreover, the other two methods are far behind (Table 1). MLE of ConvDip is lowest with 11.05*mm*, which is remarkably small considering the variance introduced by the AGM. eLORETA yields significantly larger MLE than ConvDip (*diff_median_* = 2.21*mm, p* = 10*^−^*^6^*, d* = 0.34).

**Table 1:**
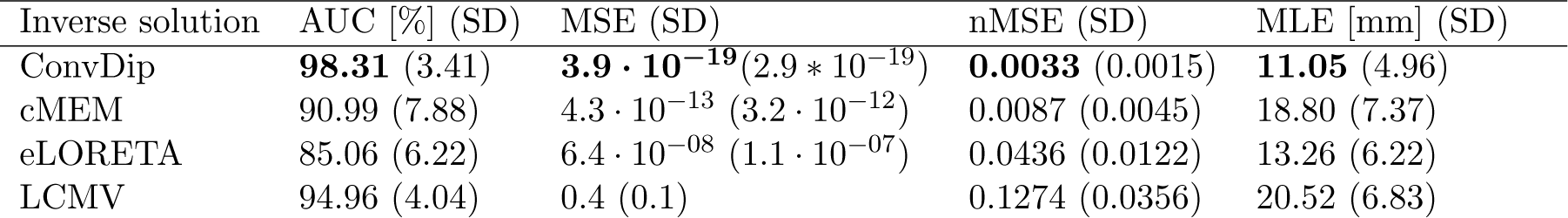
Comparison of inverse algorithm performance for samples containing a single source cluster. Source clusters were of varying spatial extents from two to seven neighborhood orders. AUC: Area under the receiver operator curve. MSE: Mean squared error, nMSE: Normalized mean squared error, MLE: Mean localization error. Best performances are highlighted in bold font.

**AUC** In Fig. 2 (left graph) the AUC for each inverse algorithm is depicted per simulated source extent. ConvDip is able to reach best performance for extents of *s*_3_ and *s*_4_ (*AUC_convdip,s_*_3_*_,s_*_4_ = 99%) with attenuated AUC especially for larger sizes (*AUC_convdip,s_*_7_ = 94%). The AUC drops by *≈* 5.2% from *s*_3_ to *s*_7_. cMEM, an inverse algorithm aiming at estimating both location and size of source clusters, shows similar properties. cMEM shows it’s best AUC at *s*_2_ (*AUC_cMEM,s_*_2_ = 93%). The AUC decreases by *≈* 5.5% from the smallest to the largest source extent, which is comparable to ConvDip. We conclude that while ConvDip yields significantly higher AUC overall (*p ≤* 10*^−^*^6^*, d >* 0.40), its behavior to varying source sizes is similar to that of cMEM. The other inverse algorithms perform significantly worse (Fig. 2)

**Figure 2:**
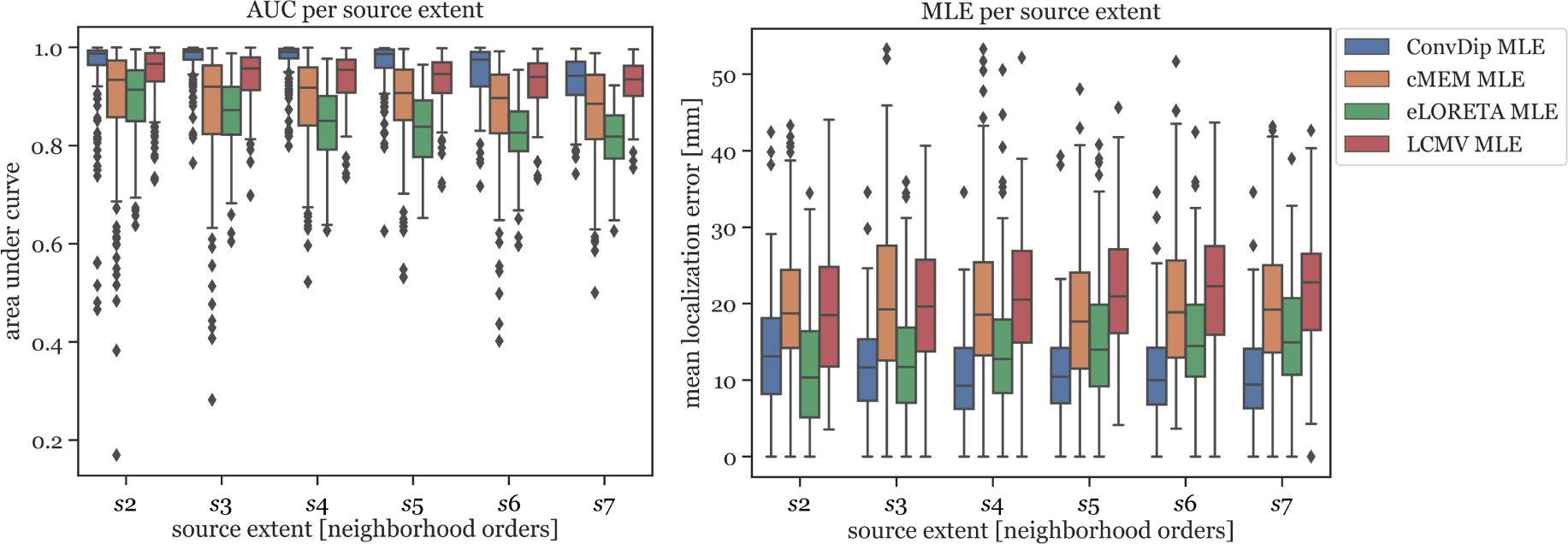
Area under the Curve and mean localization error for each inverse solution algorithm grouped by the source extent. AUC reflects capability of finding source at correct location, estimating its size and reducing false positives. MLE reflects the capability to correctly localize the center of the source cluster. Simulations used for this analysis contained only a single source cluster per sample with varying size from two to seven neighborhood orders.

**MLE** MLE of ConvDip is affected by source extent, as *MLE_convdip,s_*_2_ is 39% higher compared to *MLE_convdip,s_*_7_. This observation does not apply to the MLE yielded by inverse solutions of cMEM, which is fairly stable (7% variation). Overall, MLE was smaller for ConvDip inverse solutions compared to the other three inverse algorithms. One particular exception is the case of small source clusters (*s*_2_), for which eLORETA yields the most accurate estimates (*MLE_eLORET A,s_*_2_ = 10.34*mm*, see Fig. 2).

#### 4.1.2 Evaluation of source eccentricity

Next, we evaluated the ability of ConvDip to correctly localize source clusters of different eccentricity (i.e. distance from all electrodes). The further away a source is from the electrodes, the harder it is to localize due to volume conduction.

**AUC** In Fig. 3 (left) the AUC metric is shown for each inverse algorithm grouped by bins of eccentricity. Bins were defined from 20 mm (deep source clusters) to 80 mm (superficial source clusters). Since deeper sources are harder to localize accurately, AUC is expected to be correlated with eccentricity. However, this could be verified only for ConvDip (*r* = 0.27*, p <* 10*^−^*^16^) and cMEM (*r* = 0.31*, p <* 10*^−^*^23^). For LCMV beamformer depth is barely associated with AUC (*r* = 0.07*, p < .*05) and for eLORETA no such association could be found (*r* = 0*, p >* 0.9). Despite these interesting dynamics, ConvDip shows for all eccentricities higher AUC compared to almost all other inverse algorithms. Only the LCMV beamformer shows a non-significant tendency (*p <* 0.1*, d* = 0.2) for a higher AUC for low eccentricity of 20*mm* (*median* = 93.39% *±* 3.19) than ConvDip (*median* = 93.44% *±* 5.35).

**Figure 3:**
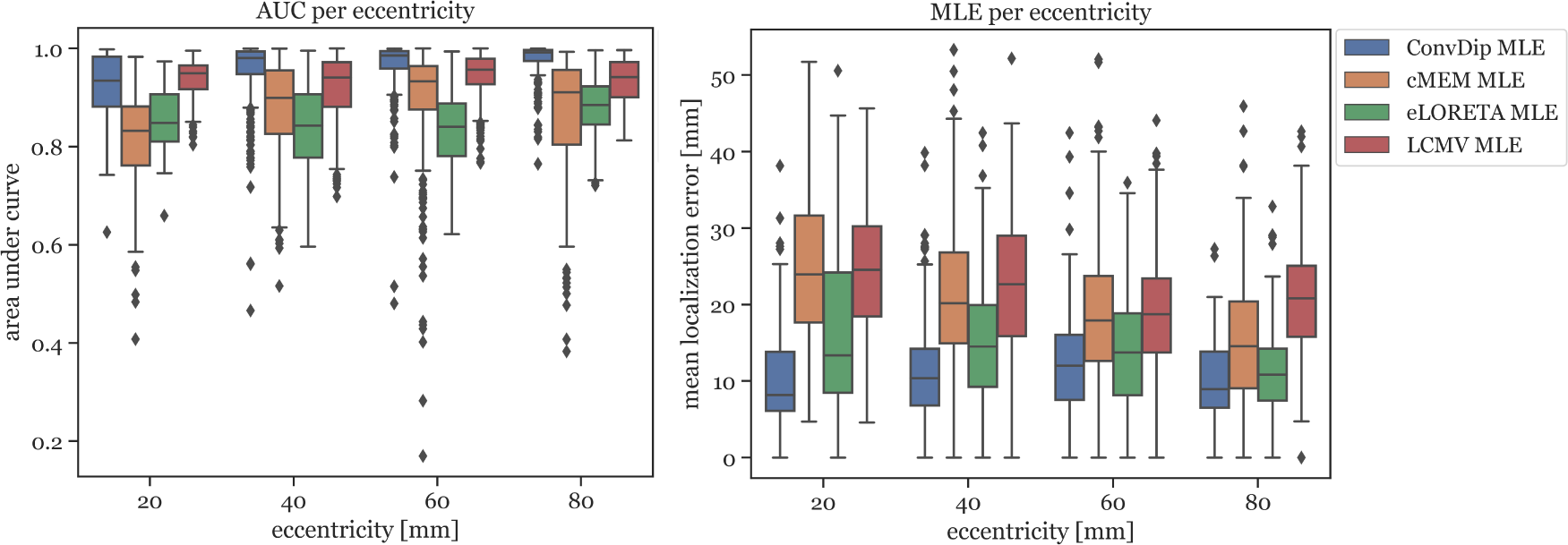
Area under the Curve and mean localization error for each inverse algorithm grouped by the source eccentricity. Simulations used for this analysis contained only a single source cluster per sample of varying size from two to seven neighborhood orders.

**MLE** MLE is also affected by source depth as can be seen in Fig. 3 (right). We calculated the ratio between the MLE of the deepest source clusters (*eccentricity* = 20*mm*) to the MLE of superficial source clusters (*eccentricity* = 80*mm*) in order to calculate depth sensitivity of each inverse algorithm, henceforth called *drop-off*. cMEM shows a drop-off of 64%, eLORETA has a drop-off of 23% and LCMV beamformer has a drop-off of 18%. ConvDip shows the lowest MLEs across eccentricities, compared to the other inverse methods. Moreover, ConvDip shows an inverted U-shaped behavior, with lowest MLE for the deepest and most superficial source clusters. MLE of ConvDip decreases slightly with depth of the source cluster and are not correlated with eccentricity (*r* = *−*0.02*, p > .*5), whereas this relation was found for all other inverse algorithms: cMEM (*r* = *−*0.30*, p <* 10*^−^*^21^), eLORETA (*r* = *−*0.15*, p <* 10*^−^*^5^) and LCMV beamformer (*r* = *−*0.09*, p <* 0.01).

Concluding, we find no drop-off in localization accuracy with depth of source clusters when using ConvDip, highlighting a systematically more robust behavior of ConvDip compared to all other inverse algorithms. With depth, however, the ability to estimate the extent of a source cluster is reduced.

#### 4.1.3 Evaluation of dipole moments

Finally, we utilized the single-source set to evaluate how well the moments of the dipoles was reconstructed by ConvDip and the other inverse algorithms. We calculated the MSE between each true and predicted source vector (as described in section 3.5). We find that the scaled predictions of ConvDip yielded the lowest MSE (3.93 *·* 10*^−^*^19^ *±* 2.91 *·* 10*^−^*^19^), followed by cMEM (4.32 *·* 10*^−^*^13^ *±* 3.24 *·* 10*^−^*^12^) and eLORETA (6.40 *·* 10*^−^*^8^ *±* 1.05 *·* 10*^−^*^7^). We excluded LCMV beamformer since the approach does not yield dipole moments that can be interpreted in this way. This result is less surprising in the light of the aforementioned results since ConvDip is able to estimate the extent of sources well, which facilitates the estimation of correct dipole moments through the scaling process (refer to section 3.3.3 for an explanation of scaling).

### 4.2 Evaluation with multiple sources

In a realistic situation the EEG may pick up signals originating from many regions in the brain simultaneously, aggravating the inversion process. Due to the low spatial resolution of the EEG not all sources can be identified reliably. In order to evaluate the performance of the inverse algorithms in reconstructing more challenging samples of the multi-source set we calculated the aforementioned metrics which are summarized in Tab. 2.

**Table 2:** Comparison of inverse algorithm performance for samples containing multiple source clusters. Samples contained between one and ten source clusters of varying spatial extent. Each cell contains the median and mean absolute deviation of the medians over all samples in the multi-source set. For the “percentage of sources found” metric the mean was calculated instead of median. AUC: Area under the receiver operator curve. nMSE: Normalized mean squared error, MLE: Mean localization error, Percentage of sources found: The percentage of sources that whose maxima was correctly localized. Best performances are highlighted in bold font.

| Inverse solution | AUC [%] (SD) | MSE (SD) | nMSE (SD) | MLE (SD) | % of sources found (SD) |
| --- | --- | --- | --- | --- | --- |
| ConvDip | **78.29** (9.39) | **$1.6 \cdot 10^{-18}$** ($4.4 \cdot 10^{-19}$) | **0.0087** (0.0042) | 28.18 (12.10) | **62.71** (22.38) |
| cMEM | 70.76 (8.95) | $1.2 \cdot 10^{-12}$ ($3.9 \cdot 10^{-12}$) | 0.0139 (0.0057) | 38.61 (16.11) | 50.12 (24.02) |
| eLORETA | 69.94 (7.54) | $1.9 \cdot 10^{-07}$ ($1.6 \cdot 10^{-07}$) | 0.0559 (0.0152) | 29.51 (8.98) | 60.71 (22.60) |
| LCMV | 71.13 (10.30) | 0.4 (0.1) | 0.1592 (0.0415) | **28.09** (5.50) | 62.36 (23.67) |

**AUC** Overall, the AUC for the multi-source set is low since the desirable 80% AUC is undercut by each inverse algorithm. ConvDip achieved the highest AUC of 78%, thereby surpassing the next best competitor LCMV by *≈* 7.2%.

**nMSE** nMSE is substantially smaller for ConvDip inverse solutions compared to all other inverse algorithms with 60% lower nMSE compared to the next best performing algorithm cMEM.

**MLE** LCMV beamforming is the only inverse solution (of the tested ones) with lower MLE than ConvDip (Table 2), albeit the absolute difference of the median MLEs is small (diff_*MLE ≈* 0.1*mm, p* = 10*^−^*^6^*, d* = 0.22).

**Percentage of found sources** The percentage of sources found was fairly similar for ConvDip, eLORETA and LCMV and *≈* 10% lower for cMEM (Table 2). Fig. 4 shows the results for the metrics AUC and nMSE grouped by the number of sources present in the samples. It is apparent that the accuracy of all inverse solutions depends strongly on the total number of sources that contribute to the EEG signal on the scalp since for each inverse algorithm, the results worsen with increasing numbers of source clusters present in the sample. Pearson correlation between AUC and the number of present source clusters was strong and statistically significant for each inverse algorithm: ConvDip (*r* = 0.70*, p <* 10*^−^*^147^), cMEM (*r* = 0.60*, p <* 10*^−^*^97^), eLORETA (*r* = 0.73*, p <* 10*^−^*^163^), LCMV (*r* = 0.73*, p <* 10*^−^*^167^).

**Figure 4:**
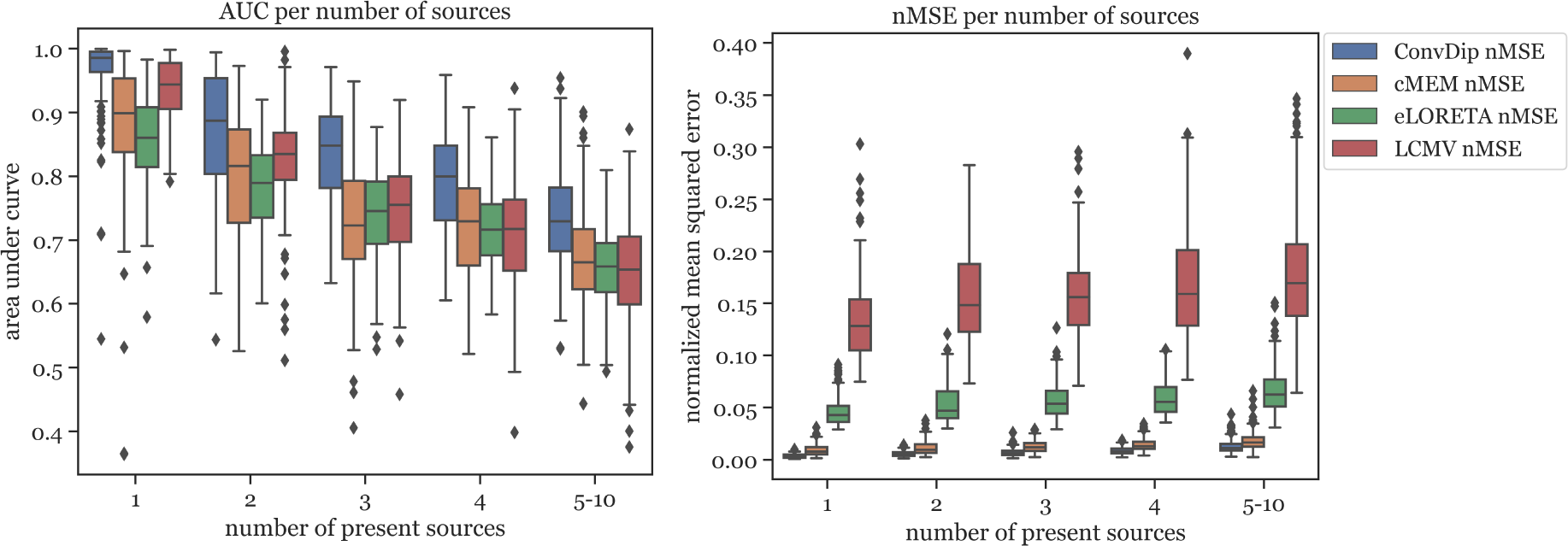
Area under the Curve and normalized mean squared error for each inverse algorithm grouped by the number of present source clusters. The right-most columns in each of the graphs display the results for samples containing between five and ten source clusters.

**Figure 5:**
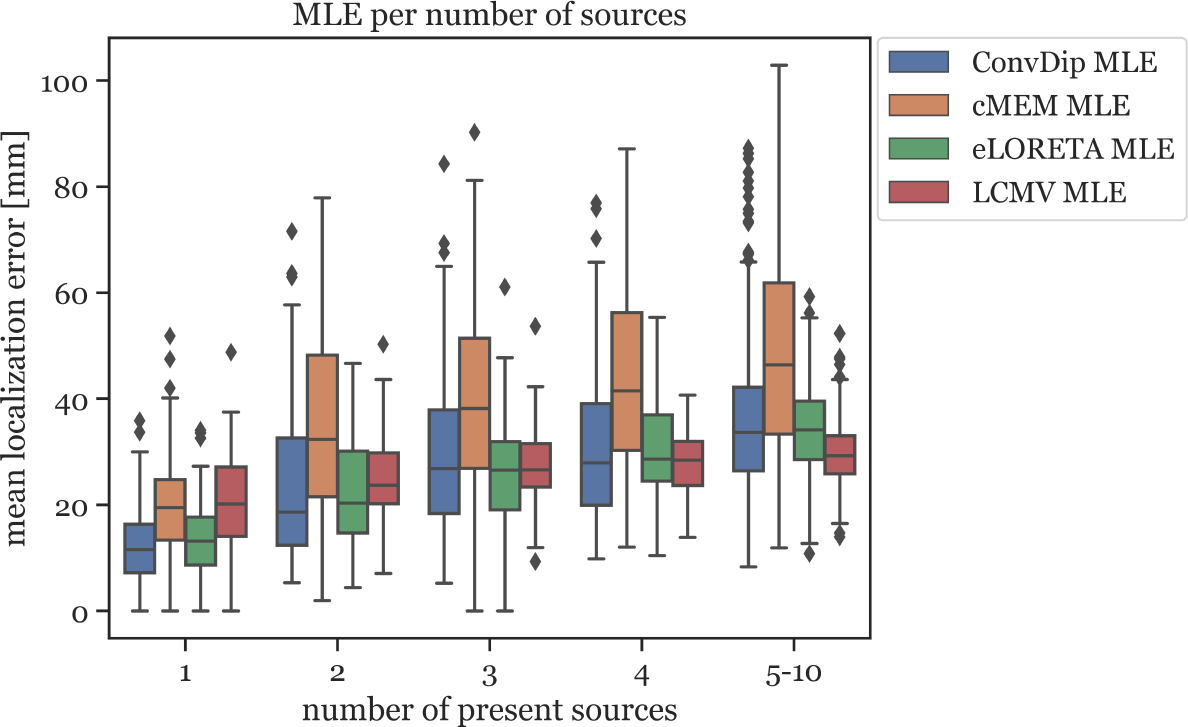
Mean localization error for each inverse algorithm grouped by the number of present source clusters. The rightmost column displays the results for samples containing between five and ten source clusters.

This relationship was also found to exists between MLE and the number of source clusters. For large numbers of source clusters (*≥* 5), LCMV shows the lowest MLE (29.41*mm*) or 4.24*mm* less than ConvDip (*p* = 10^6^*, d* = 0.60). This may be a result of the inherent noise-suppressing properties of the LCMV beamformer.

A core result of this analysis is that although ConvDip was trained with samples containing between one and five source clusters it is capable of inverting more challenging samples as well (Fig. 4).

Furthermore, we showed that discrepancies between the assumed generative model and the true bio-physical properties underlying an EEG measurement (e.g. tissue conductivity, precise electrode locations, spacing of the discrete dipole model) can be handled by ConvDip at least to some degree.

ConvDip provides inverse solutions that are valid for samples generated with the AGM and different configurations (namely, different numbers of source clusters). This shows, that ConvDip is able to generalize well beyond samples within the training set. This in turn confirms the validity of this purely data-driven approach to handle the EEG inverse problem.

## 5 Discussion

### 5.1 Overview

We have presented ConvDip, a convolutional neural network that aims to find extended sources of EEG signals. The approach of using a neural network to solve the inverse problem is distinct from classical methods, mainly because it works purely data-driven. All constraints on the solution of ConvDip are given implicitly. The performance of ConvDip was evaluated and tested against commonly used inverse algorithms cMEM, eLORETA and LCMV beamforming. In the following we discuss the significance of these results.

### 5.2 Using ANNs to solve the inverse problem

One major result of this work is that the CNN ConvDip was capable to reliably find the location of neural sources that give rise to the EEG. ConvDip transforms a 7 *×* 11 image to a large vector of 5124 voxels. In image classification, CNNs are utilized to do the opposite: To extract hidden information from a large input matrix and output a single value that indicates some property, e.g. the presence of a cat or a dog. Our work verifies that CNNs are capable to solve an underdetermined problem. This issue was already addressed by Lucas, Iliadis, Molina, and Katsaggelos (2018), who demonstrated the advantages of ANNs over analytical approaches to solve the inverse problem.

We trained ConvDip on data generated by GM and subsequently evaluated ConvDip on simulated samples that were generated by the AGM in order to avoid committing the inverse crime. Overall, the performance of ConvDip was sometimes comparable but in most cases better than the classical inverse solutions cMEM, eLORETA and LCMV beamformer. This provides evidence that ConvDip learned to generalize beyond samples in the training data and remains robust to deviations from the training data. This holds for small deviations in the position of the electrodes (*±*3.7*mm*), tissue conductivities, but also in the configuration of sources (number and size of clusters, deviations from the GM source model grid) and provides another remarkable advantage of this data driven approach.

We evaluated the performance of ConvDip and the alternative inverse algorithms concerning multiple objectives, the results of which we will discuss in the following.

### 5.3 Estimating source extent

The ability to estimate not only the location of a neural source but also its extent was tested for all inverse algorithms on the single-source set using the AUC and nMSE performance. We find that ConvDip reaches a nearly perfect AUC of *≈* 98% which is not met by any other inverse algorithm. nMSE, a measure of global similarity between the true and estimated source, was smallest using ConvDip, with less than half the error compared to the next best inverse solutions yielded from cMEM (Tab. 1). ConvDip furthermore estimated different source sizes appropriately (Fig. 2) with higher AUC than all other inverse algorithms. Although these findings are remarkable we have to regard that the shape and size of the sources was rather predictable for ConvDip. All sources in the training and evaluation set had a more or less circular shape due the region growing approach we chose. It may be presumably a large challenge for ConvDip to estimate ellipitical shapes or shapes that are even more deviant to a circle. Despite this caveat, it is remarkable how well ConvDip performs in this comparison considering the variability introduced by the AGM.

### 5.4 Localizing deep sources

A further surprising result for us during the evaluation was that for ConvDip the MLE did not depend on the depth of the source cluster, i.e. ConvDip is capable to correctly localize even deep sources with nearly no drop-off (see Fig. 3). This finding was a strong contrast to the findings from other inverse algorithms, which all tend to mislocate deeper sources more than superficial ones. Although source depth did not influence the MLE of ConvDip we find that the AUC is in deed compromised by depth. This holds also true for cMEM but not for eLORETA and barely for LCMV beamformer. Overall, ConvDip achieved comparable AUCs and MLEs for various depth of source clusters, compared to other inverse solutions, rendering it a viable option for localizing deep sources.

### 5.5 Performance with multiple active source clusters

In subsection 4.2 we evaluated how ConvDip performs when many source clusters are simultaneously active. We observed a drop-off in AUC with increasing numbers of active source clusters from *≈* 99% with single sources to *≈* 73% when 5 or more sources were active simultaneously. Similar behavior was observed for all inverse algorithms tested (see Fig. 4). But still, ConvDip achieved the highest AUC compared to all alternatives tested, surpassing the best competitor LCMV by *≈* 7.2%. Furthermore, ConvDip also outperformed all other inverse algorithms concerning nMSE (Tab. 2). Only when five or more source clusters are present we find slightly lower MLE using LCMV beamformer compared to ConvDip. This increased localization ability of LCMV beamformer could be due to the active suppression of noise signals in the beamforming process. Another reason could be that ConvDip was never trained with samples that contained more than five active source clusters, therefore struggling to accurately localize at least a subset of the source clusters.

### 5.6 Computation speed

As already pointed out by Sclabassi, Sonmez, and Sun (2001), ANNs naturally yield faster inverse solutions than classical methods. On our workstation (CPU: Intel i5 6400, GPU: Nvidia RTX 2070, 16 Gb RAM) one forward pass of ConvDip took on average *≈* 31*ms* on the GPU. When all necessary preprocessing steps (such as re-referencing, scaling and interpolation to 2-D scalp maps) are taken into account we reached *≈* 32*ms* of computation time on average. In comparison, it takes *≈* 129*ms* (*≈*factor 4) to calculate the eLORETA inverse solution using MNE, *≈* 98*ms* ms (*≈*factor 3) for LCMV beamforming and astonishing 11.55*s* (*≈*factor 360) to calculate the iterative cMEM inverse solution using brainstorm. Additionally one has to provide at the noise covariance matrix, which takes on average *≈* 45*ms* to compute using the empirical estimation of MNE. In practical terms ConvDip is capable to compute 31.25 inverse solutions per second (ips) on a GPU, whereas eLORETA reaches 7.75*ips* LCMV beamformer 10.20*ips* on a CPU. Possibly, implementations of eLORETA and LCMV beamforming that can be run on a GPU achieve lower computation times compared to CPU. The short computation times of ConvDip are at a distinct advantage over other inverse solutions, e.g. in real-time applications as in neurofeedback experiments or in the development of brain-computer interfaces. When comparing our approach to the existing literature on ANN-based inverse solutions, we can now confirm that an ANN not only provides competitively accurate single dipole locations but also a distributed dipole model of the brain-electric activity, taking into account more than one dipole. This can be attributed mostly to the rapid developments in the machine learning domain such as the introduction of convolutional layers (LeCun & Bengio, 1995) and the improvements of graphical processing units that render the training of large ANNs possible in acceptable time.

### 5.7 Realistic simulations

The simulation of EEG data in this study was based on different assumptions on the number, distribution and shape of electromagnetic fields; inspired by Nunez and Srinivasan (2006). The validity of predictions of any inverse solution is only granted if the assumptions on the origin of the EEG signal are correct. This is indeed one of the critical challenges towards realistic inverse solutions in general, both for data-driven inverse solutions (such as ConvDip) as well as classical inverse solutions with physiological constraints (e.g. eLORETA). Therefore, when interpreting the applicability of ConvDip to real data, it is important to specify and justify these assumptions. We show that the knowledge about brain-electric activity underlying EEG signals is emulated by ConvDip after it was trained. However, it is evident from the present work, how critical the particular choice of parameters is with respect to the performance, as ConvDip produces solutions that closely resemble the training samples (see Appendix A, Appendix B). A more sophisticated approach to this problem would be to diversify the training data, e.g. by changing the region-growing procedure to produce sources with higher variety of shapes beyond simple spheric source clusters. Nonetheless, ConvDip finds reasonable source clusters when presented with real data (see Appendix C for an example).

An important observation in real EEG data is the inter-individual variability, which is in part a direct result of individual brain anatomies. For ConvDip application to real EEG data, particularly with group data, it is thus advised to collect anatomical MRI brain scans of each individual subject. Individual neural networks then need to be trained based on each subjects’ individual anatomy provided by the MRI data. This may help to achieve accurate group-study source estimations. This, however, raises the problem of computation time, since training individual CNNs is time-consuming. A solution to this is an inter-individual transfer learning approach, where ConvDip is trained on one subjects’ anatomy and fine-tuned for each additional subject with new training data of the individual anatomies. Fine tuning could be achieved by replacing the output-layer, lowering the learning rate and retraining for only few epochs. Training time *per se* is another important topic, when considering CNNs to handle the inverse problem.

### 5.8 Training time

Although the availability of high-performing hardware resources (e.g. high performing graphic cards) is growing rapidly, the training of the presented neural network architecture requires a considerable amount of processing power and processing time. The version of ConvDip shown in this work required *≈* 10.5*hours* of training for 500 epochs and 100,000 samples of data using a NVIDIA RTX 2070 graphical processing unit. Improving the architecture of ConvDip to shorten training time is an important task for future research, especially when individualized models are required. Limiting the solution space to fewer voxels may be one way to decrease the computation time since most trainable parameters are located at the final fully-connected layers. The easiest way would be to reduce the number of parameters of the neural network, e.g. by targeting a CNN with volumetric output of the ANN corresponding to a volumetric source model of the brain. This could spare the computationally expensive fully-connected layers and reduce the number of weights in the network. Another possibility is to segment the cortical surface into parcels, i.e. clusters of neighboring dipole positions. This would reduce the size of the output layer by one to two orders of magnitude and thereby reduce the number of trainable parameters dramatically. The feasibility of such a dimension reduction is to be investigated in future developments.

Another computationally expensive processing step is the generation of realistic artificial training data. In this study, we projected neural activity using a three-layer BEM head model. Generating 100,000 simulations of neural activity took 12 minutes using a PC with CPU @ 4×4.3GHz and 16 GB of RAM. Software that enables EEG researchers to perform physiologically realistic simulations of neural activity for their own ConvDip application is available from different resources, e.g. from the MNE library in Python (Gramfort et al., 2014).

### 5.9 Further perspectives for improvement

The present version of ConvDip is trained on single time instances of EEG data. From a single EEG data point it is, however, not possible to estimate a noise baseline. As a consequence, such a baseline has not yet been taken into account in ConvDip, which leaves room for improvements towards more regularized and adaptive solutions that are computed flexibly depending on noise conditions. In contrast, LCMV, eLORETA and cMEM explicitly make use of a noise covariance matrix which is used to regularize the inverse solution. ConvDip could be improved by extending the data input to an EEG time series, allowing the neural network to learn spatio-temporal patterns.

Possible further advances could be made by preserving the 3D structure of the output space by, instead of a flattened output layer. This can be realized using a deconvolutional neural network for 2D-3D projections (Lin, Kong, & Lucey, 2018) or by using spatiotemporal information with a LSTM network (Cui et al., 2019). One obvious further way to improve the performance of ConvDip may be to increase the capacity of the neural network, e.g. by adding more layers. In our initial testing phase we tried different neural architectures without observing significant improvements by adding more layers that justify the increase in training time. Neural architecture search should be used to find a compact, well-performing architecture to solve the EEG inverse problem (Liu et al., 2018).

Finally, the development of a both meaningful and computationally inexpensive loss function which makes training faster and allows for faster convergence is key for ANN-based approaches to the inverse problem that can run on ordinary PC within a reasonable amount of time.

### 5.10 Outlook

We showed that ConvDip, a shallow CNN, is capable of solving the inverse problem using a distributed dipole solution. The association between single time points of EEG measurements and underlying source distributions can be learned and used to predict plausible inverse solutions. These inverse solutions were furthermore shown to globally reflect the topology of the brain-electric activity given by the training data.

Predictions yielded from ConvDip are in rare cases comparable to but in the majority of cases better than the here tested existing inverse algorithms in the measures we focused on. Further, it the application of ConvDip to real EEG data yields reasonable results. The fully trained ConvDip requires only little computational costs of 32 ms, which makes it a promising tool for real-time applications where only single time points of EEG data are available.

We want to emphasize the importance of realistic simulations of *real* neural activity that can be measured by EEG. Although a lot of knowledge about the generation of scalp potentials was implemented in our EEG simulations, including more physiological constraints can further reduce the complexity of the inverse problem. One example is the estimation of cortical column orientation (Bonaiuto et al., 2019) or the incorporation of empirical knowledge about region-specific activity from in-vivo (e.g. electrocorticogram) studies. Taking into account these additional aspects may further improve ConvDip’s performance.

The current ConvDip version was based on single time points of artificial EEG data. Exploiting temporal aspects of brain dynamics (neighboring time points) may provide additional valuable information and increase the predictive power that may further refine future ConvDip versions.

## 6 Appendix A

**Figure 6:**
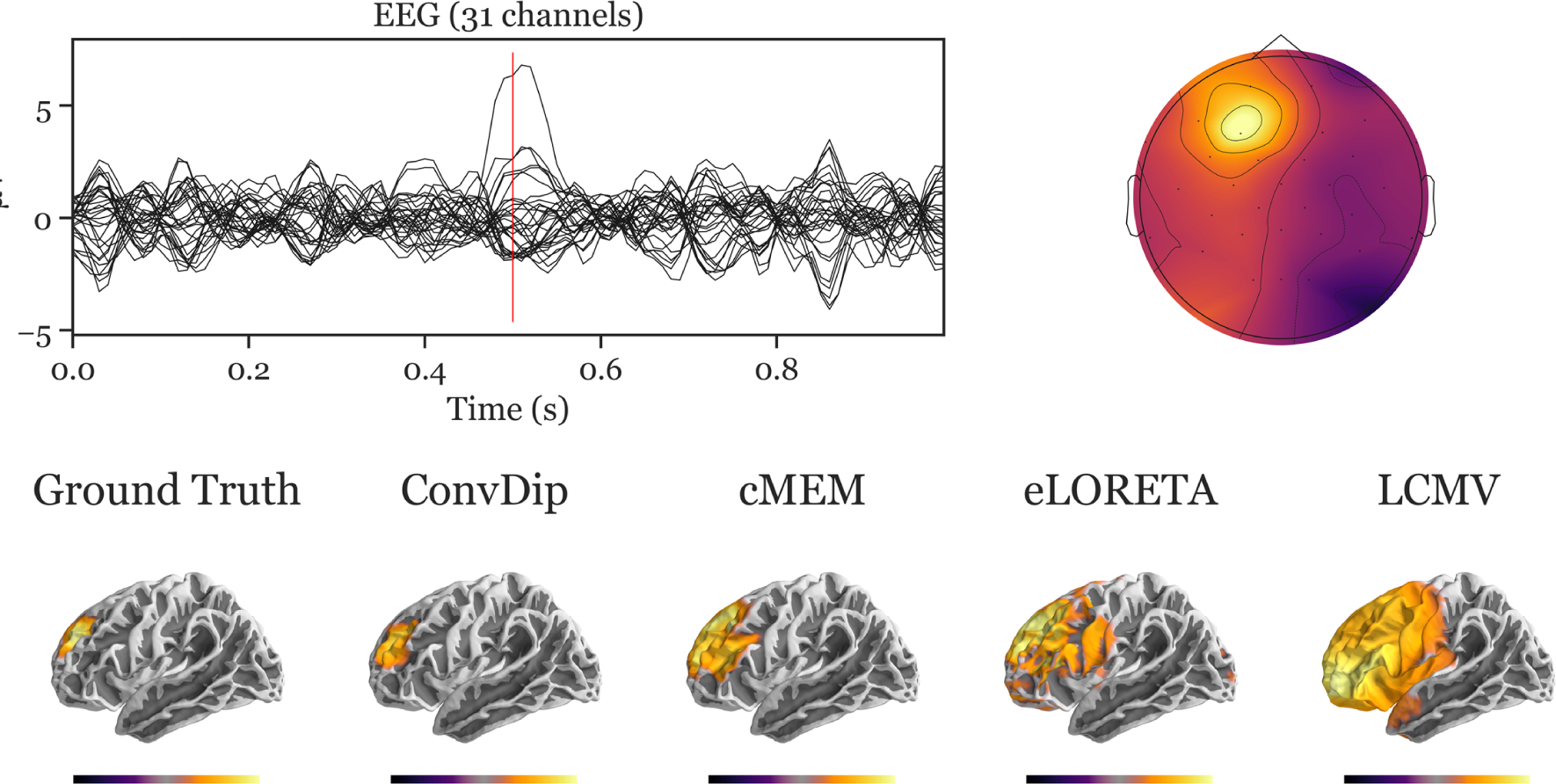
Inverse solution of a simulation containing a single source cluster. An exemplary, representative sample of inverse solutions for a single source cluster. (A) depicts the ERP at each of the 31 channels containing both signal (central peak) and realistic noise from real recordings. (B) shows the scalp map at the central ERP peak (as indicated by the vertical red line in A). (C) shows the dipole moments plotted on the white matter surface of the template brain in lateral view of the left hemisphere. On the *left* the ground truth source pattern is depicted with a source cluster in the frontal cortex of the left hemisphere. Various inverse solutions that aim to recover this pattern are depicted next to it. Voxels below 25% of the respective maximum are omitted for a clearer representation of the current distribution.

## 7 Appendix B

**Figure 7:**
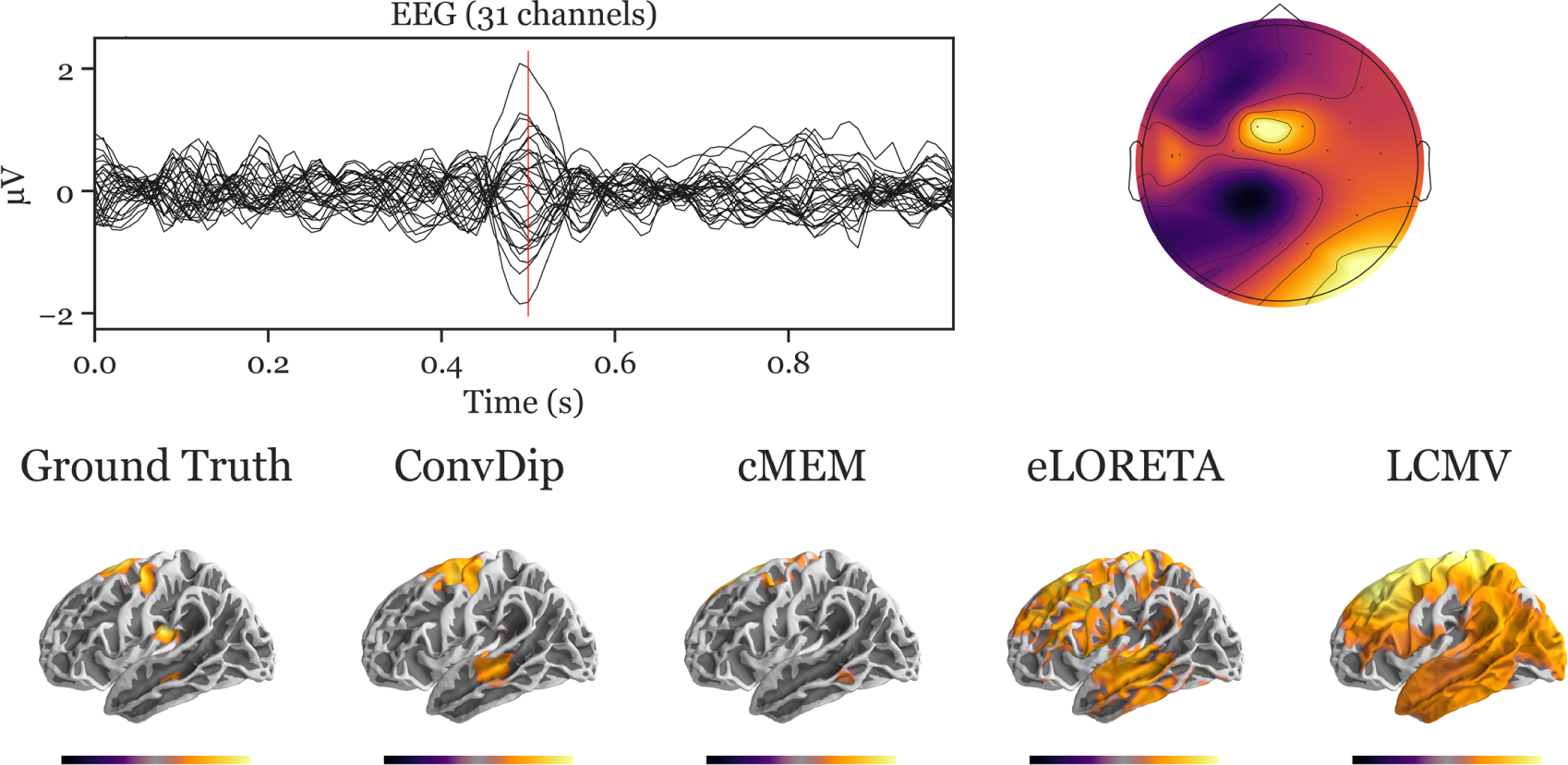
Inverse solution of a simulation containing four source clusters. An exemplary, representative sample of inverse solutions for four source clusters. (A) depicts the ERP at each of the 31 channels containing both signal (central peak) and realistic noise from real recordings. (B) shows the scalp map at the central ERP peak (vertical red line in A). (C) shows the dipole moments plotted on the white matter surface of the template brain in lateral view of the left hemisphere. On the *left* the ground truth source pattern is depicted with a source cluster in the motor cortex, supplementary motor area, insula and the middle temporal lobe of the left hemisphere. Various inverse solutions that aim to recover this pattern are depicted next to it. Voxels below 25% of the respective maximum are omitted for a clearer representation of the current distribution.

## 8 Appendix C

**Figure 8:**
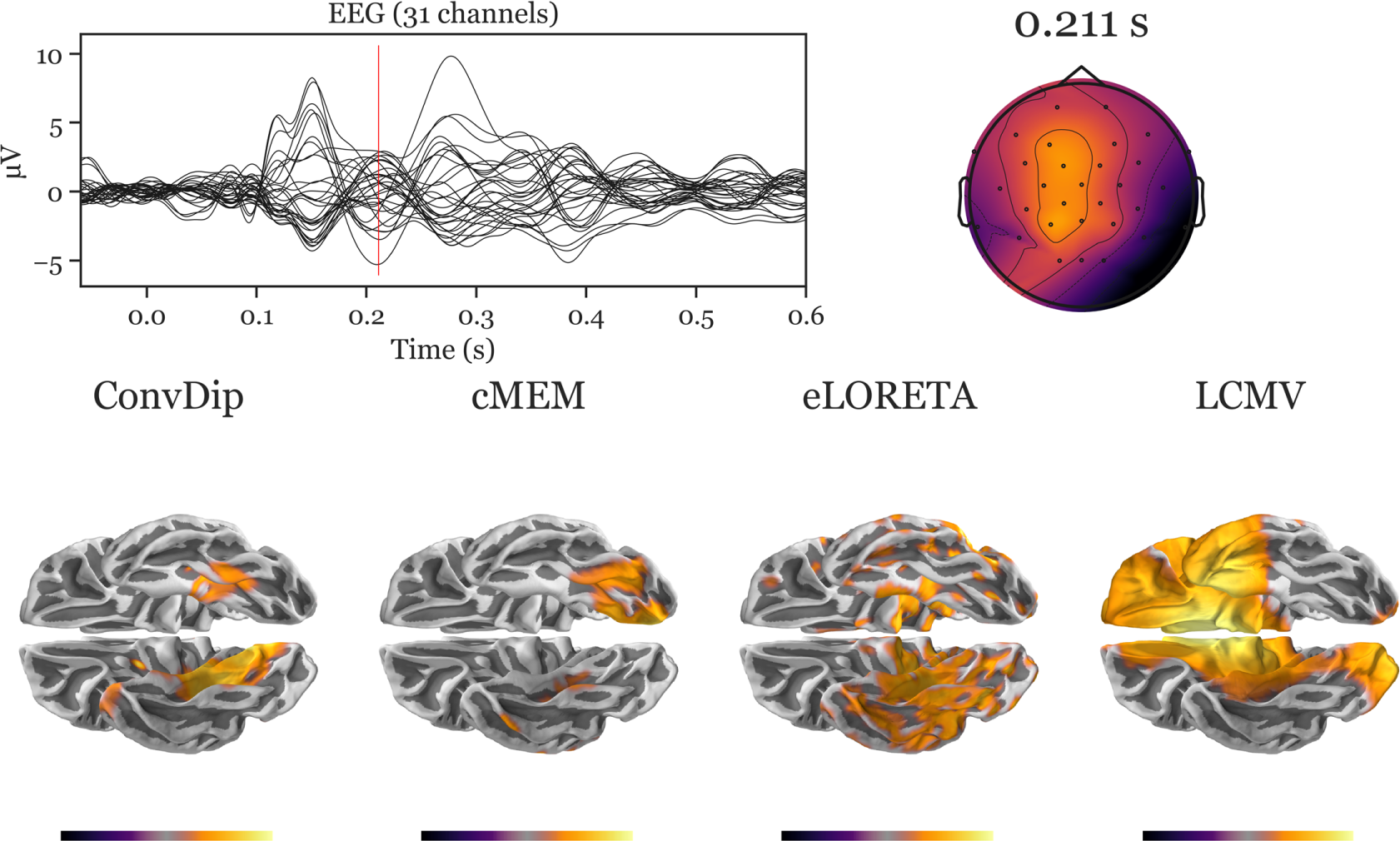
Inverse solution of real face-evoked data. All inverse algorithms were subjected to real EEG data recorded from a single participant containing 150 trials of face-evoked potentials. (A) depicts the ERP at each of the 31 channels. (B) shows the scalp map at a N170-like component (vertical red line in A, 211 ms after stimulus onset). (C) shows the dipole moments plotted on the white matter surface of the template brain in ventral view of the right hemisphere to reveal the inferior temporal cortex (ITC). Various inverse solutions show activity in the ITC. Notably, ConvDip and cMEM recover a focal source cluster close or within the fusiform face area, a region known to be involved in face processing (e.g. Joos et al., 2020; Kanwisher & Yovel, 2006). Voxels below 25% of the respective maximum are omitted for a clearer representation of the current distribution.

## Footnotes

^1^Implementation was adapted from https://github.com/N0vel/weighted-hausdorff-distance-tensorflow-keras-loss/

